# Complete mitochondrial genomes of riverine *Lamprologus* (Actinopterygii, Cichlidae) with an emphasis on the blind cichlid *L. lethops*

**DOI:** 10.1101/2024.09.11.612419

**Authors:** Sebastian M. Jimenez, Naoko P. Kurata, Melanie L. J. Stiassny, S. Elizabeth Alter, Prosanta Chakrabarty, Fernando Alda

## Abstract

Lamprologine cichlids are a diverse group of fishes distributed in Lake Tanganyika and the Congo River. Nine species of *Lamprologus* occur in the Congo River basin including the only blind cichlid *Lamprologus lethops*, but little is known about the natural history and evolution of this enigmatic species. To alleviate this knowledge gap, we characterized the complete mitochondrial genomes of *L. lethops* and its riverine congeners and provided a phylogenetic hypothesis based on these data. We recovered complete mitochondrial genomes from eleven specimens of eight species of *Lamprologus*. Mitogenomes were identical in the number and order of genes and similar in size (16,579-16,587 bp). In contrast to previous phylogenomic studies, riverine *Lamprologus* were recovered in two non-sister mitochondrial lineages that were more closely related to other genera of Lake Tanganyika lamprologines than to each other. In the first lineage, most relationships were not highly supported. In the second lineage, *L. lethops* was recovered as the sister species of *L. markerti, L. mocquardi* and *L. tigripictilis*. Interestingly, sequences from *L. mocquardi* were found in the two mitochondrial lineages. Our results hint at multiple events of past introgression and highlight the importance of increasing taxonomic and genomic sampling to study complex evolutionary histories.

## 1. INTRODUCTION

The genus *Lamprologus* comprises 20 species within the tribe Lamprologini that are mainly found in Lake Tanganyika and associated streams but unlike other genera within the lamprologines, *Lamprologus* also includes fully riverine species found throughout the Congo River drainage (Ronco et al., 2020; Schelly and Stiassny, 2004; Stiassny and Alter, 2021). Riverine taxa exhibit unique adaptations to the extreme environmental conditions they encounter (Kurata et al., 2022; Stiassny and Alter, 2021). The most remarkable species in the lower Congo River (LCR) is *Lamprologus lethops* Roberts & Stewart, 1976, the only known cryptophthalmic and fully depigmented cichlid species (Fig. 1).

**Fig. 1.**
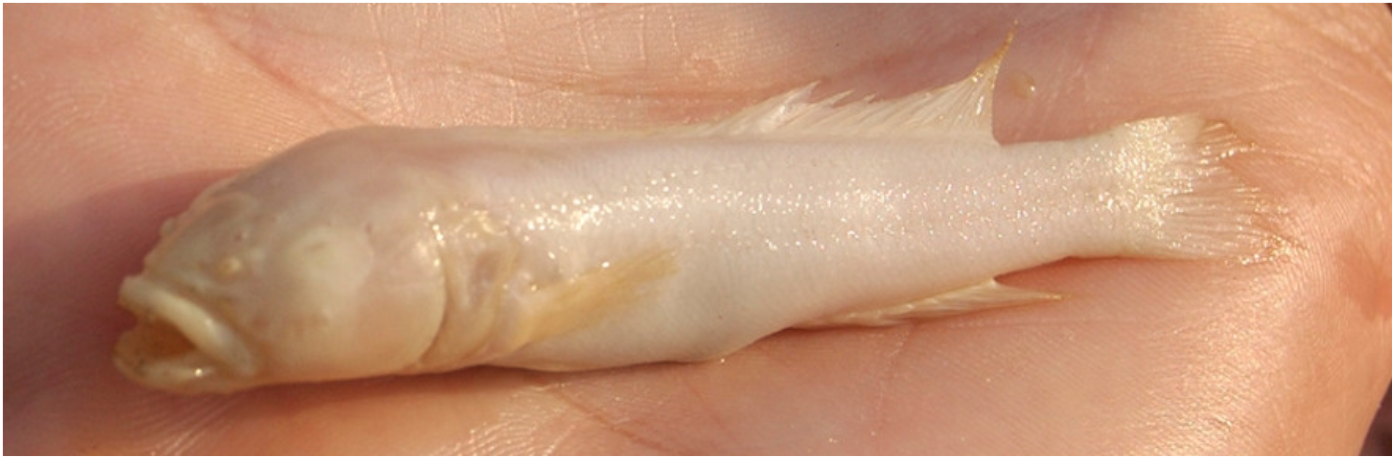
Specimen of *L. lethops* (c. 70 mm long) on a hand. Photograph by the American Museum of Natural History, used with permission and courtesy of Melanie L. J. Stiassny.

Recent genome-wide phylogenetic analyses support the monophyly of the riverine *Lamprologus* species, although high levels of gene tree discordance preclude confident resolution of the relationships within the riverine clade (Astudillo-Clavijo et al., 2022; Alda et al. in preparation). Mitochondrial DNA studies do not support the monophyly of this group, but these studies are often incomplete in their taxonomic or in their gene sampling (Day et al., 2007; Schedel et al., 2019). Here, we aim to contribute to the resolution of these knowledge gaps by presenting the first report of complete mitochondrial genomes for eight riverine species of *Lamprologus* including the blind cichlid *L. lethops*, and a preliminary mitogenomic hypothesis.

## 2. MATERIALS AND METHODS

We analyzed 11 samples of eight species of *Lamprologus* that inhabit the Congo River: *Lamprologus congoensis* Schilthuis, 1891 (AMNH 257860, AMNH255211), *Lamprologus lethops* Roberts & Stewart, 1976 (AMNH263957), *Lamprologus markerti* Tougas & Stiassny, 2014 (AMNH238650), *Lamprologus mocquardi* Pellegrin, 1903 (ZSM Kis-2008-080, ZSM Kis-2008-003), *Lamprologus teugelsi* Schelly & Stiassny, 2004 (AMNH238649), *Lamprologus tigripictilis* Schelly & Stiassny, 2004 (AMNH263989), *Lamprologus tumbanus* Boulenger, 1899 (AMNH247886), and *Lamprologus werneri* Poll, 1959 (AMNH239703, AMNH263499) (Table 1). We also analyzed two lamprologine species: *Julidochromis dickfeldi* Staeck, 1975 (AMNH26257) and *Telmatochromis burgeoni* Poll, 1942 (AMNH264405), and sequences from available partial mitogenomes (Schedel et al., 2019), that were used as outgroups (Table 1). All fishes except *L. lethops* were collected alive and euthanized in accordance with ethical and legal guidelines for international animal research approved by the AMNH Institutional Animal Care and Use Committee (IACUC) (approval #36/06). Specimens and tissues are deposited in the Ichthyology collections of the American Museum of Natural History (https://nhm.org/research-collections/departments-and-programs/ichthyology, curator: Melanie L. J. Stiassny) and the Zoologische Staatssammlung München (Bavarian State Collection of Zoology; https://zsm.snsb.de/sektion/ichthyology/?lang=en, curator: Ulrich Schliewen) (Location and voucher information in Table 1).

**Table 1.**
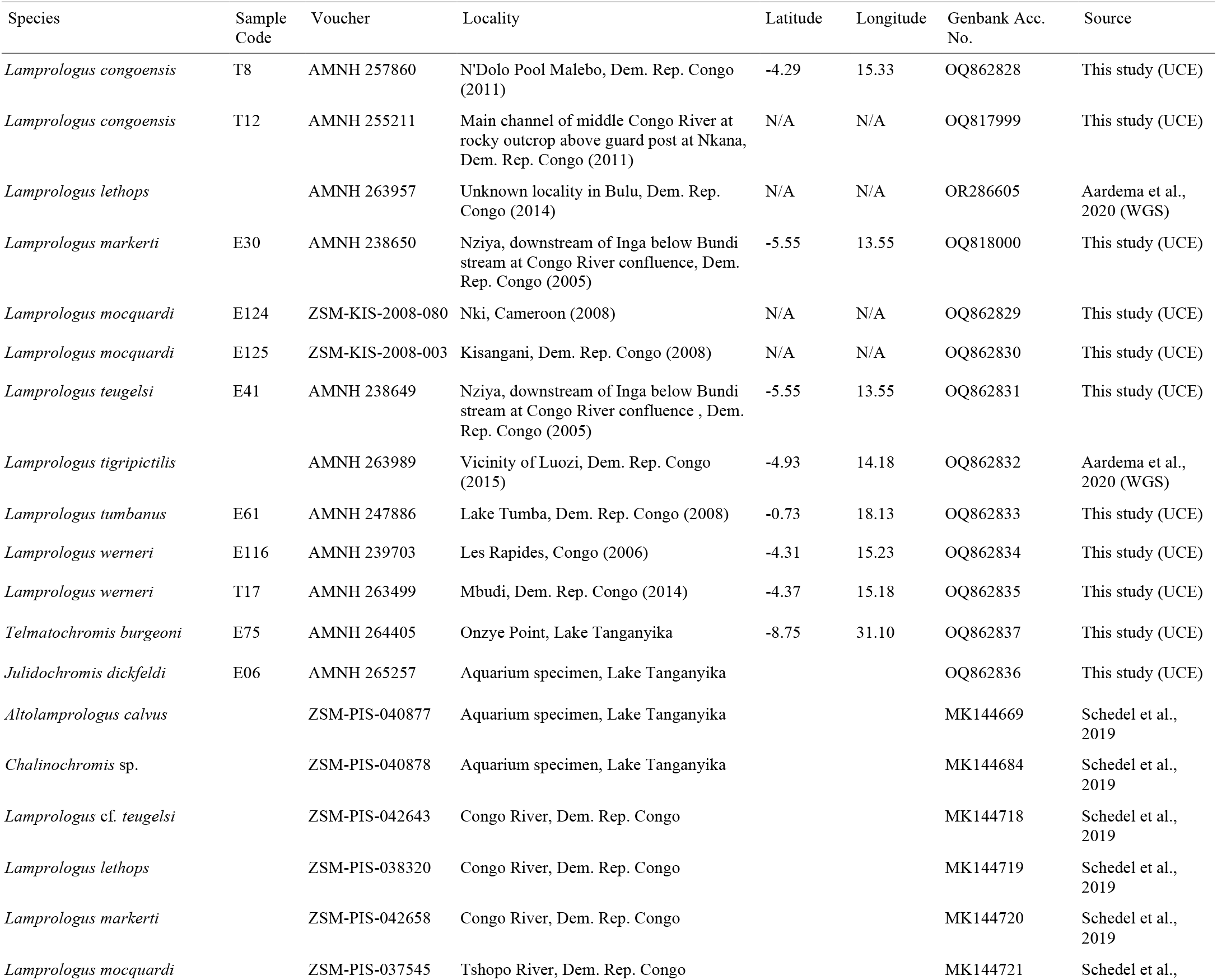
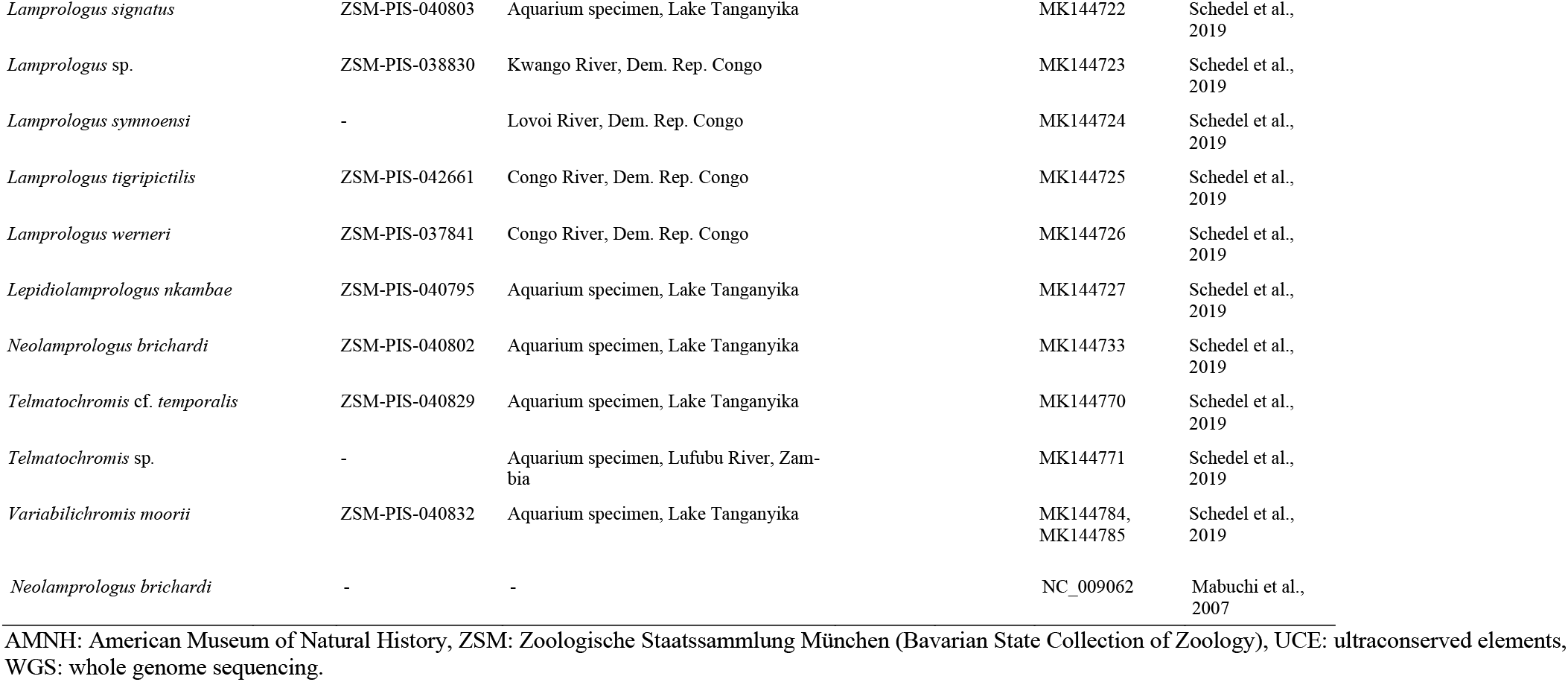
List of samples analyzed in this study.

We obtained mitochondrial sequences as a by-product of hybrid target capture of ultraconserved elements (UCEs) (PRJNA1097814) and whole genome libraries (Aardema et al., 2020, BioProject PRJNA577474). We used Geneious Prime 2022.1.1 (https://www.geneious.com) to trim our raw reads for low-quality bases (cut-off limit: 0.05) and map them to the closest reference mitochondrial genome available (*Neolamprologus brichardi*, NC_009062, Mabuchi et al., 2007).

We extracted all protein-coding genes from our complete mitochondrial genomes and aligned them with the partial genomes available using MAFFT 1.5.0. (Katoh and Standley, 2013). We partitioned the data by gene and codon and estimated the best arrangement of partitions and their nucleotide substitution model using Partition Finder (Lanfear et al., 2016) in IQ-TREE2 (Minh et al., 2013). We inferred a maximum-likelihood tree and assessed nodal support using 1000 ultrafast bootstrap replicates (Hoang et al., 2018). Finally, we used the approximately unbiased (AU) test (Shimodaira, 2002) to test the monophyly of the riverine *Lamprologus* by comparing an unconstrained maximum-likelihood tree and a constrained tree where riverine species form a monophyletic group.

## 3. RESULTS

### 3.1. Characteristics of *Lamprologus* mitogenomes

We recovered complete genomes from all individuals except for *L. markerti*, which was missing 33 bp in the *ND3* gene and 93 bp in the *ND5* gene, and *L. teugelsi* which was missing 174 bp in the *ND1* gene. The mean coverage of the genomes recovered from UCE raw sequence data was 57.96✕ ±18.78 SD, and 3872✕ ±702.10 SD in those recovered from whole genome sequencing data (Fig. S1). All species genomes consisted of 22 tRNA genes, two rRNA genes, 13 protein-coding genes, and a control region (D-loop) in identical order. The mean base composition was 26.95% ±0.121 SD of A, 30.94% ±2.2682 SD of C, 16.31% ±0.197 SD of G, and 26.67% ±0.205 SD of T. The genome lengths ranged from 16,579 bp in *L. lethops* to 16,587 bp in *L. mocquardi* (Fig. 2).

**Fig. 2.**
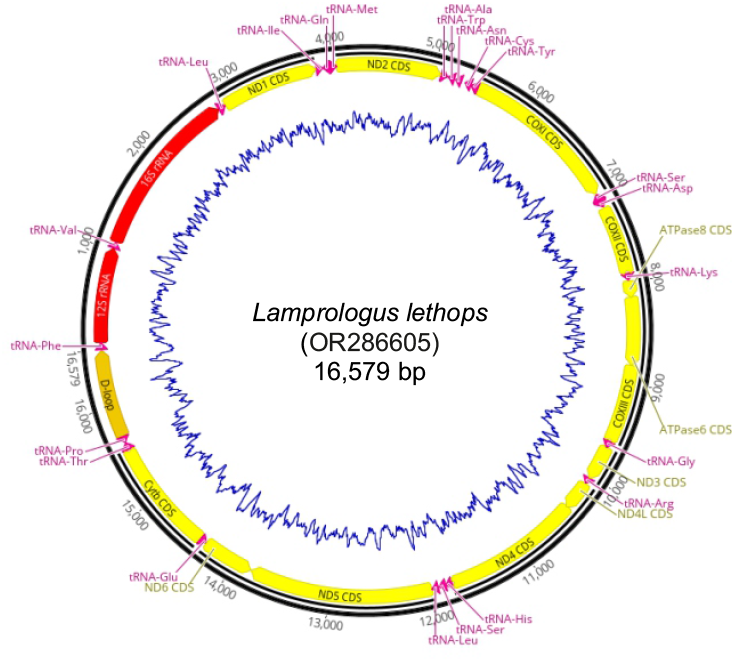
Graphical representation of the complete mitochondrial genome of *L. lethops* showing the annotation of all protein-coding genes (yellow arrows), rRNA genes (red arrows), tRNA genes (pink arrows), and control region (tan arrow). The blue line represents GC content. For color codes, refer to the online version of this article.

The most common start codon was ATG (Met), except for GTG (Val) that was the start codon in the *COX1* gene in all species. The most common stop codon was TAA followed by T--. In *ND1*, TAG was the stop codon in all species except in *L. teugelsi* that used TAA. The stop codon of *ND2* was T--in *L. lethops, L. markerti, L. teugelsi*, and *L. tigripictilis*, and TA-in *L. congoensis, L. mocquardi, L. tumbanus*, and *L. werneri* (Table S1).

### 3.2. Mitochondrial phylogeny of riverine *Lamprologus*

The best maximum-likelihood tree recovered species of riverine *Lamprologus* in two lineages (Fig. 3). The first lineage included a clade composed exclusively of riverine *Lamprologus* (*L. werneri, L. congoensis, L. mocquardi, L. tumbanus*). The second lineage also included a clade of riverine *Lamprologus*, in which *L. markerti* and *L. mocquardi* were closely related and sister to *L. tigripictilis*, and *L. lethops* was sister to them. *Lamprologus teugelsi* and *L*. sp. from the Kwango River also belonged to this clade (Fig. 3).

**Fig. 3.**
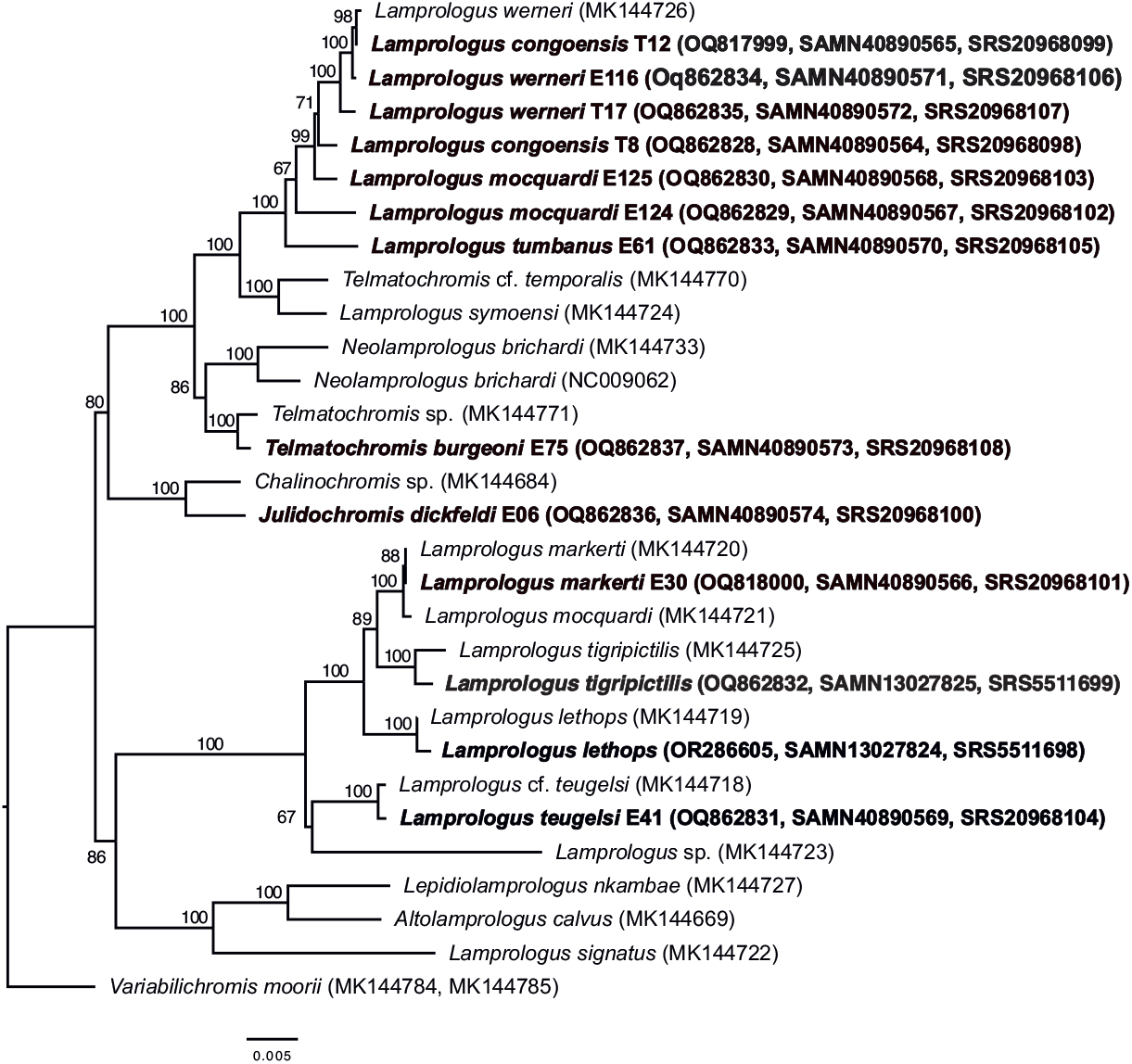
Maximum-likelihood (IQ-TREE 2) phylogeny using all mitochondrial protein-coding genes of the riverine *Lamprologus* species analyzed in this study. Numbers next to nodes are support values obtained after 1000 ultrafast bootstrap replicates, and numbers between parentheses are GenBank accession numbers, BioSample numbers, and SRA numbers. The following sequences were obtained in this study and are highlighted in bold: OQ817999, OQ818000, OQ862828, OQ862829, OQ862830, OQ862831, OQ862832, OQ862833, OQ862834, OQ862835, OQ862836, OQ862837 and OR286605. The following sequences were obtained from the literature: NC_009062 (Mabuchi et al. 2007), and MK144669, MK144684, MK144718, MK144719, MK144720, MK144721, MK144722, MK144723, MK144724, MK144725, MK144726, MK144727, MK144733, MK144770, MK144771, MK144784*, MK144785* (Schedel et al., 2019). *Note that *Variabilichromis moori* is a combination of two partial mitochondrial genome sequences. Sample details are indicated in Table 1.

## 4. DISCUSSION AND CONCLUSION

Mitogenomes of riverine *Lamprologus* were identical in the number and order of genes and similar in size to other lamprologines (Mabuchi et al., 2007). We recovered the riverine *Lamprologus* in two non-sister mitochondrial lineages, in which individuals of the same species (*L. congoensis, L. mocquardi, L. werneri*) were not always resolved as each other’s closest relatives.

Genetic sequence data from the enigmatic *L. lethops* are scarce and ours is the first complete mitochondrial genome for the species (Fig. 2). Similar to recent genomic studies, mitogenomic data recovered *L. lethops* in a clade including LCR species such as *L. tigripictilis, L. markerti, L. werneri* and *L. teugelsi* (Astudillo-Clavijo et al., 2022). Despite the differences, all studies suggest that *L. lethops* originated through a process of in situ speciation within the LCR.

Schedel et al. (2019) had recovered a clade exclusively formed by LCR species. However, when we added more specimens to this study, the monophyly of the LCR group was rejected (p-value of AU test = 1.32 × 10^−7^). This finding contrasts with morphological and phylogenomic hypotheses that strongly support the monophyly of riverine *Lamprologus* (Astudillo-Clavijo et al., 2022; Schelly, 2006; Schelly and Stiassny, 2004). Mito-nuclear discordance and even complete mitochondrial replacements are not uncommon among syntopic lamprologines in Lake Tanganyika (Koblmüller et al., 2007; Nevado et al., 2009; Schelly, 2006) and both past and ongoing introgressive hybridization may explain this pattern. In the Congo River, on the other hand, *Lamprologus* species show allopatric or parapatric distributions, with little gene flow inferred between neighboring species (Kurata et al., 2022), thus it seems probable that ancient hybridization events explain the discordance observed.

Overall, our data suggest multiple hybridization events, uncover undescribed mitochondrial lineages and highlight the necessity of analyzing larger sample sizes and multiple genomic compartments to fully investigate processes of reticulated evolution in this fish group.

## Acknowledgements

We thank the Honors College and Brock Scholar Program from the University of Tennessee at Chattanooga for providing support for S.M.J. For assistance with collection and exportation permitting M.L.J.S. is grateful to the Ministère de l’Agriculture et du Développement Rural, Secrétariat Géneral de l’Agriculture, Pêche et Elevage, Direction des Pêches and the Université de Kinshasa, Cabinet du Recteur, in the Democratic Republic of Congo.

## CREDIT AUTHORSHIP CONTRIBUTION STATEMENT

SMJ and FA conceived the project, NPK, MLJS, SEA, and FA did the experimentation and obtained molecular data, SMJ carried out formal analyses with supervision from FA, MLJS, SEA, PC and FA provided samples and other resources, MLJS identified the samples, SMJ and FA wrote the first draft, and NPK, MLJS, SEA and PC validated, reviewed and edited the final manuscript.

## BIOGRAPHICAL NOTE

SMJ is a B.S. General Biology undergraduate student at the University of Tennessee at Chattanooga and participated in this study as part of a funded project from the UTC Honors College. SMJ has participated in an NSF summer REU and eventually, intends to go on to graduate school. NPK is currently a postdoctoral research associate in the Department of Natural Resources and the Environment at Cornell University. NPK’s research uses genomic and paleoclimatic data to investigate drivers of diversification, and adaptive evolution of fishes. MLJS is a research curator in the Department of Ichthyology of the American Museum of Natural History. MLJS’s research focuses on the systematics and evolutionary morphology of tropical freshwater fish faunas. Ongoing projects are centered in describing fish diversity in central Africa and elucidating the mechanisms of species diversification. SEA is an Associate Professor at California State University Monterey Bay. SEA’s research focuses on evolutionary and ecological genetics of aquatic species drawing from evolutionary genetics, phylogenomics, predictive habitat modeling and isotope studies. PC is a Professor and Fish Curator at the Louisiana State University Museum of Natural Science. PC is a systematist and an ichthyologist studying the evolution and biogeography of both freshwater and marine fishes. FA is currently a Staff Scientist at the Spanish National Research Council (CSIC). FA’s research investigates the relative roles of ecological and geological processes in driving genetic differentiation and speciation.

## ETHICAL APPROVAL

Work was conducted following the Institutional Animal Care and Use Committee (IACUC) protocols provided by the American Museum of Natural History (approval #36/06).

## DATA AVAILABILITY STATEMENT

The raw sequence data that support the findings of this study are openly available in GenBank of NCBI at https://www.ncbi.nlm.nih.gov/ under BioProjects PRJNA577474 (BioSample numbers: SAMN13027824 and SAMN13027825, SRA numbers: SRX6989555 and SRX6989556) and PRJNA1097814 (BioSample numbers: SAMN40890564, SAMN40890565, SAMN40890566, SAMN40890567, SAMN40890568, SAMN40890569, SAMN40890570, SAMN40890571, SAMN40890572, SAMN40890573 and SAMN40890574; and SRA numbers: SRX24194295, SRX24194296, SRX24194297, SRX24194298, SRX24194299, SRX24194300, SRX24194301, SRX24194302, SRX24194303, SRX24194304, SRX24194305). The accession numbers of the assembled mitochondrial genomes are: OQ817999, OQ818000, OQ862828, OQ862829, OQ862830, OQ862831, OQ862832, OQ862833, OQ862834, OQ862835, OQ862836, OQ862837 and OR286605 (see Table 1 and Fig. 3 for details). Additional supporting data for this study (sequence alignments and annotated genomes as General Feature Format files) are openly available in Zenodo at https://doi.org/10.5281/zenodo.11442789.

## SUPPLEMENTARY MATERIALS

**Fig. S1** Read depth of coverage plot of all the mitochondrial genomes assembled in this study. For reference, grey boxes indicate the location of the mitochondrial protein coding genes.

**Table S1** Characteristics of the mitochondrial genomes of the *Lamprologus* species analyzed in this study. The first and last nucleotide positions (bp) are shown for *L. lethops* and the differences in size relative to this specimen are shown for the rest of species (rel. size). The start and stop codons are indicated for each coding gene.

