## Supplementary Material for "Complete mitochondrial genomes of riverine *Lamprologus* (Actinopterygii, Cichlidae) with an emphasis on the blind cichlid *L. lethops*"

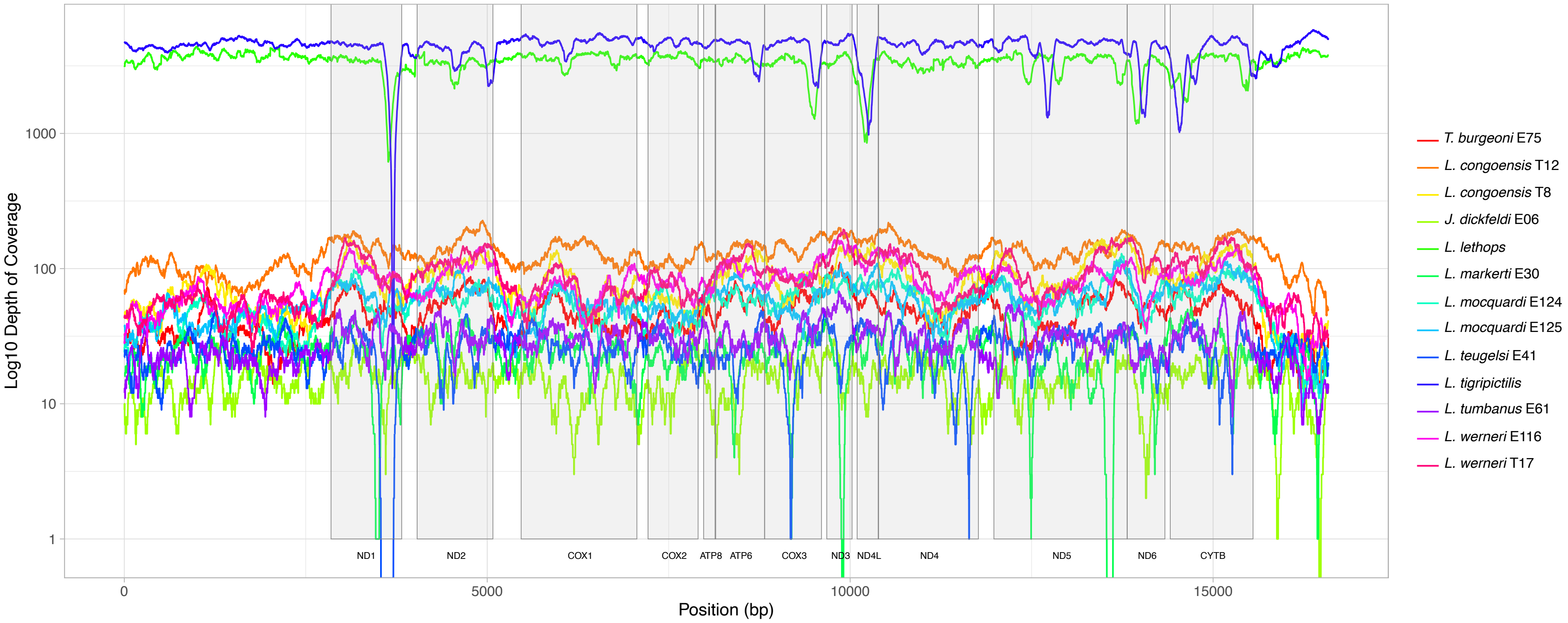


**Fig. S1** Read depth of coverage plot of all the mitochondrial genomes assembled in this study. For reference, grey boxes indicate the location of the mitochondrial protein coding genes.
